# The role of phages in proglacial stream biofilm communities characterized by microdiversity

**DOI:** 10.1101/2023.09.25.559239

**Authors:** Hannes Peter, Gregoire Michoud, Susheel Bhanu Busi, Tom J. Battin

## Abstract

Viruses modulate the diversity and activity of microbial communities. However, little is known about their role for the structure of stream biofilms communities. Here, we present insights into the diversity and composition of viral communities in various streams draining three proglacial floodplains in Switzerland. Proglacial streams are characterized by extreme environmental conditions, including near-freezing temperatures and ultra-oligotrophy. These conditions select for few but well-adapted bacterial clades, which dominate biofilm communities and occupy niches via microdiversification. We used metagenomic sequencing to reveal a remarkably diverse biofilm virome in these streams. Across the different floodplains and streams, viral community composition was tightly coupled to that of the bacterial hosts, which was underscored by generally high host specificity. Combining predictions of phage-host interactions with auxiliary metabolic genes (AMGs), we identify specific AMGs shared by phages infecting microdiverse clade members. Our work provides a step towards a better understanding of the complex interactions among bacteria and phages in stream biofilm communities in general and streams influenced by glacier meltwaters and characterized by microdiversity in particular.

## Introduction

Viruses infecting bacteria, also called bacteriophages, play important roles in modulating the diversity and activity of bacterial assemblages in every biome on Earth (Dion et al., 2020). Particularly phage-induced mortality (Suttle, 1994, 2007) is known to control bacterial abundance and biomass, community composition and diversity with sizable impacts on global biogeochemical cycles (Roux et al., 2016; Zimmerman et al., 2020). However, beyond mortality, phages also influence the fitness of their bacterial hosts in numerous ways. Lysogenic infections, for example, can impede host functioning by insertion of phage genetic material into functional genes (Obeng et al., 2016). Phage-encoded sigma factors can influence bacterial spore formation, affecting important bacterial life-history traits such as dormancy (Schwartz et al., 2022). Moreover, metabolic reprogramming during lytic infections can scale to ecosystem-scale biogeochemical consequences (Zimmerman et al., 2020).

While the consequences of viral infections for individual bacterial cells are often detrimental, beneficial effects of phage-host interactions typically arise at the level of populations, communities or even holobionts (Shkoporov et al., 2022). For instance, phage-driven diversification (Koskella and Brockhurst, 2014), lysogenic conversion (Howard-Varona et al., 2017) or auxiliary metabolic genes (AMGs) can increase phenotypic plasticity or augment the metabolic repertoire of microbial assemblages (Shkoporov et al., 2022). In biofilms, where densely packed communities of microbes interact, phage genomic material has been shown to enter even phylogenetically distant hosts directly or during conjugative transfers, ultimately increasing the immunological memory (Hwang et al., 2023).

Besides the density, diversity and activity of phages and their hosts, the eco-evolutionary consequences of phage infections depend on host specificity. While early studies suggested predominantly narrow phage host ranges, more recent work, including single-cell and metagenomics studies of hydrothermal biofilms (Jarett et al., 2020; Hwang et al., 2023), suggest indeed a continuum of host ranges (de Jonge et al., 2019).

Here, we present metagenomic insights into the interactions between bacterial biofilm communities and their phages in high-altitude stream ecosystems. Alpine streams are extreme environments, characterized by ultra-oligotrophy, low water temperature, high UV exposure and short ice-free seasons, yet represent hotspots of microbial biodiversity (Freimann et al., 2013; Hotaling et al., 2017; Brandani et al., 2022; Busi et al., 2022; Fodelianakis et al., 2022).

Glacier influence adds additional constraints for life in alpine streams. Close to the glacier terminus, glacier-fed streams (GFS) are permanently ice cold and the erosive activity of glaciers liberates large quantities of mineral particles, which render GFS turbid and limit light for phototrophic primary production. Compared to GFS, groundwater-fed streams (GWS) that drain elevated terrain towards the edges of the floodplains, are warmer, their streambeds more consolidated and lower turbidity allowing photosynthetic primary producers to dwell (Freimann et al., 2013; Brandani et al., 2022).

Previous work on biofilm communities in GFS and GWS revealed that they are astonishingly diverse (Brandani et al., 2022), distinct from the microbial community suspended in the water column (Ezzat et al., 2022) and shaped predominantly by environmental selection (Fodelianakis et al., 2022; Brandani et al., 2023). More specifically, this means that ecologically successful taxa in both GFS and GWS are recruited from similar clades, including taxa classified as *Polaromonas*, *Rhodoferax*, *Rhizobacter*, *Methylotenera* and *Massilia*. These clades are microdiverse (Fodelianakis et al., 2022), indicating that they efficiently fill available niches which cannot be occupied by less-well adapted taxa. Microdiversity is an intrinsic property of many microbial communities (Larkin and Martiny, 2017), including marine viral communities (Needham et al., 2017; Gregory et al., 2019). By focusing on microdiverse biofilm communities in proglacial streams, we aim to shed new light on the role of host-phage interactions in microbial communities. We hypothesize that narrow phage-host ranges predominate in communities dominated by microdiverse clades because selective constraints which lead to microdiversification may also shape bacteria-phage co-evolution. Moreover, we speculate that interactions between phages and microdiverse clade members in extreme environments may be predominantly beneficial, thus contributing to the ecological success of these clades in extreme environments. We argue that studying microbial communities in alpine stream biofilms is ideally suited to address such questions, because of the pronounced microdiversity and limited dispersal due to geographic and topographic isolation, which can increase phage-host encounter probabilities and hence lessen the co-evolutionary strength of phage-host associations.

## Material and Methods

### Study sites

We here present data from an extensive survey of benthic biofilm communities sampled from proglacial streams in the Swiss Alps. Detailed descriptions of the sampling design and environmental parameters as well as analyses of spatial diversity patterns (Brandani et al., 2022), assembly processes (Brandani et al., 2023) and the functional diversity (Michoud et al., 2023) are available elsewhere.

Briefly, we collected 47 biofilm samples from the coarse sandy sediment fraction (0.25–3.15 mm) from GFS (n=12) and GWS (n=35) from the Otemma Glacier (OTE, 45.95E, 7.45N), the Valsorey Glacier (SOY, 45.91E, 7.27N) and the Val Roseg Glacier (VAR, 46.39E, 9.84N) floodplains. Sediment samples were collected using flame-sterilized equipment, immediately flash-frozen on dry ice and stored at -80°C until processing. Samples for bacterial cell counting were directly fixed using paraformaldehyde/glutaraldehyde, flash frozen and kept at-80°C. After extraction from sediments using pyrophosphate, shaking and sonication, cells were stained using SybrGreen and counted using flow cytometry (NovoCyte, ACEA)(Brandani et al., 2022; Fodelianakis et al., 2022).

### DNA extraction, library preparation, and sequencing

DNA was extracted from 0.5 g of sediment using an optimized extraction protocol tailored to the low-biomass and mineral nature of these samples(Busi et al., 2020). Metabarcoding libraries were prepared using 2-3 ng µL^-1^ input DNA and primers targeting the V3-V4 hypervariable region of the 16S rRNA gene (341f (5’-CCTACGGGNGGCWGCAG-3’) and 785r (5’-GACTACHVGGGTATCTAATCC-3’)). Amplification was performed on a Biometra Trio (Biometra) using the KAPA HiFi DNA Polymerase (Hot Start and Ready Mix formulation) in a 25 μL-amplification reaction containing 1 x PCR buffer, 1 μM of each primer, 0.48 μg μL^−1^ BSA and 1.0 μL of template DNA. After an initial denaturation step at 95°C for 3min, 25 cycles of 94°C for 30 s, 55°C for 30 s and 72°C for 30 s were followed by a final extension at 72°C for 5 min. Sequencing libraries were prepared using dual indices (Illumina). Prior to paired-end sequencing on a single MiSeq (Illumina) lane, library DNA concentrations were quantified, normalized and pooled. Libraries were then sequenced at the Lausanne Genomic Technologies Facility (Switzerland).

Metagenome libraries were prepared using the NEBNext Ultra II FS library kit, with 50 ng of input DNA. Briefly, DNA was enzymatically fragmented for 12.5 min and amplified using 6 PCR cycles. Qubit (Invitrogen) and Bioanalyzer (Agilent) were used to check the quality and insert size of the libraries (450 bp). The metagenomes were sequenced at the Functional Genomics Centre Zürich using an S4 flowcell on a NovaSeq (Illumina) platform.

### Bioinformatic processing

Amplicon sequences were quality checked using Trimmomatic v0.36 (Bolger et al., 2014) and assigned to exact Amplicon Sequence Variants (ASVs) using DADA2 (Callahan et al., 2016) implemented in QIIME2 (Bolyen et al., 2019) with default parameters. Singleton ASVs were removed and taxonomy was assigned using the naïve Bayesian classifier implemented in QIIME2 and the SILVA v138.1 reference database. Non-bacterial ASVs (i.e. chloroplasts, mitochondria and archaea) were discarded. We built a phylogenetic tree using RAxML (Stamatakis, 2014) and the GTRCAT substitution model and the rapid bootstrapping option.

Metagenomic sequence reads were preprocessed using trim_galore (Krueger et al., 2021) v0.6.6, which uses fastqc v0.11.9 and cutadapt v3.4 for quality control and adapter removal. Trimmed reads were assembled with megahit v.1.2.9 (Li et al., 2015) using default parameters and a minimum contig length of 1,000 bp. Prokaryotic Metagenome-Assembled Genomes (MAGs) were generated *de novo* using MetaBAT2 v2.15 (Kang et al., 2019) and concoct v1.1 (Alneberg et al., 2014). Reads were mapped to the assemblies with CoverM v0.6.1. DAS Tool v1.1.2 (Sieber et al., 2018) was used to obtain a non-redundant set of MAGs for each sample. All MAGs were then dereplicated with dRep v3.2.2 (Olm et al., 2017), considering a completeness level greater than 75% and a contamination level less than 10% as determined by checkM v1.1.3 (Parks et al., 2015). Taxonomy of MAGs was assigned using GTDB-Tk v1.7.0 (Chaumeil et al., 2019). DefenseFinder v1.0.9 (Tesson et al., 2022) using default settings was used to identify antiviral systems on MAGs.

Putative viral contigs were identified from the assemblies using VIBRANT v1.2.1 (Kieft et al., 2020) with default parameters. VIBRANT also provides information about auxiliary metabolic genes on viral contigs. Complete viral contigs were then identified using CheckV v0.8.1 (Nayfach et al., 2021) and set aside. The remaining, non-complete viral contigs were binned into vMAGs with PHAMB v1.0.1 (Johansen et al., 2022) which uses VAMB v3.0.2(Nissen et al., 2021) and DeepVirFinder v1.0 (Ren et al., 2020) to obtain high-quality viral genomes.

Subsequently, complete viral genomes and high-quality vMAGs were dereplicated using vRhyme v1.1.0 (Kieft et al., 2022) taking the longest bin and 97% identity to obtain a final set of vMAGs. CheckV was then used to assess vMAG completeness and quality. BACPHLIP (Hockenberry and Wilke, 2021) was used to assign temperate or virulent lifestyles to complete vMAGs. Viral taxonomy was assigned using kaiju v.1.9.0 (Menzel et al., 2016) and the *viruses* database (only viruses from the NCBI RefSeq database). PHIST v1.1.0 (Zielezinski et al., 2022), an alignment-free, k-mer frequency based tool was used to predict phage-host interactions and iPHoP v1.3.2 (Roux et al., 2023), a consensus-approach host identification tool was used to validate these interactions.

### Statistical analyses

The statistical software language R was used to prepare figures and to perform all statistical analyses. Specifically, viral taxonomic composition was visualized using the heat_tree function of the *metacoder* (Foster et al., 2017) R package. Multivariate analyses of bacterial and viral community composition included non-metric multidimensional scaling ordination (*metaMDS*), procrustes correlation (*procrustes* and *protest*), and permutational multivariate analysis of variance using distance matrices (PERMANOVA, *adonis2*) were performed using the *vegan* (Oksanen et al., 2013) R package. Prior to these analyses, Wisconsin double standardization was applied to relative abundances and AMG counts were Hellinger transformed prior to Principal Component Analysis. Phage-host interactions were visualized using Cytoscape v 3.10.0 (Shannon et al., 2003).

## Results and Discussion

### Bacterial microdiversity in proglacial streams

In total, we obtained 35,170 bacterial ASVs across the three different floodplains and the different stream types. The bacterial communities were taxonomically dominated by members of Gamma- and Alphaproteobacteria (17.4 and 8.2% of ASVs, respectively), Bacteroidia (12.5%), Planctomycetes (7.8%) and Verrucomicrobiae (6.4%). This diversity was further classified into 925 different genera, of which members of *Candidatus* Nomurabacteria (1.8% of ASVs), *Flavobacterium* (1.7%), and *Candidatus* Saccharimonadales (1.4%) were the most prevalent ones.

Previous work has identified environmental factors common to both GFS and GWS, such as low water temperature and ultra-oligotrophy to shape proglacial stream biofilm communities (Brandani et al., 2022). Yet, important environmental differences between GFS and GWS select for different ASVs, nested within these genera and accounting for the compositional (Freimann et al., 2013; Brandani et al., 2022, 2023) and functional differences (Michoud et al., 2023) between stream types. Here, we used phylogenetic distances between ASVs to identify microdiverse bacterial clades. *Polaromonas*, *Rhodoferax*, *Rhizobacter*, *Methylotenera* and *Massilia* are particularly prevalent in proglacial streams and featured consistently short phylogenetic distances among ASVs (Figure 1a). Short phylogenetic distances could be a consequence of the number of ASVs with the same taxonomic classification. Indeed, we found that genera with few ASVs had generally shorter phylogenetic distances than genera containing many ASVs (Figure 1b). *Polaromonas*, *Rhodoferax*, *Rhizobacter*, *Methylotenera* and *Massilia,* however, contain hundreds of ASVs, comparable to other, non-microdiverse bacterial genera. And while these five genera account on average (±SD) for only 5.0±2.3% of ASV richness, they contribute 14.5±11.3% to relative abundance (Figure 1c,d).

**Figure 1.**
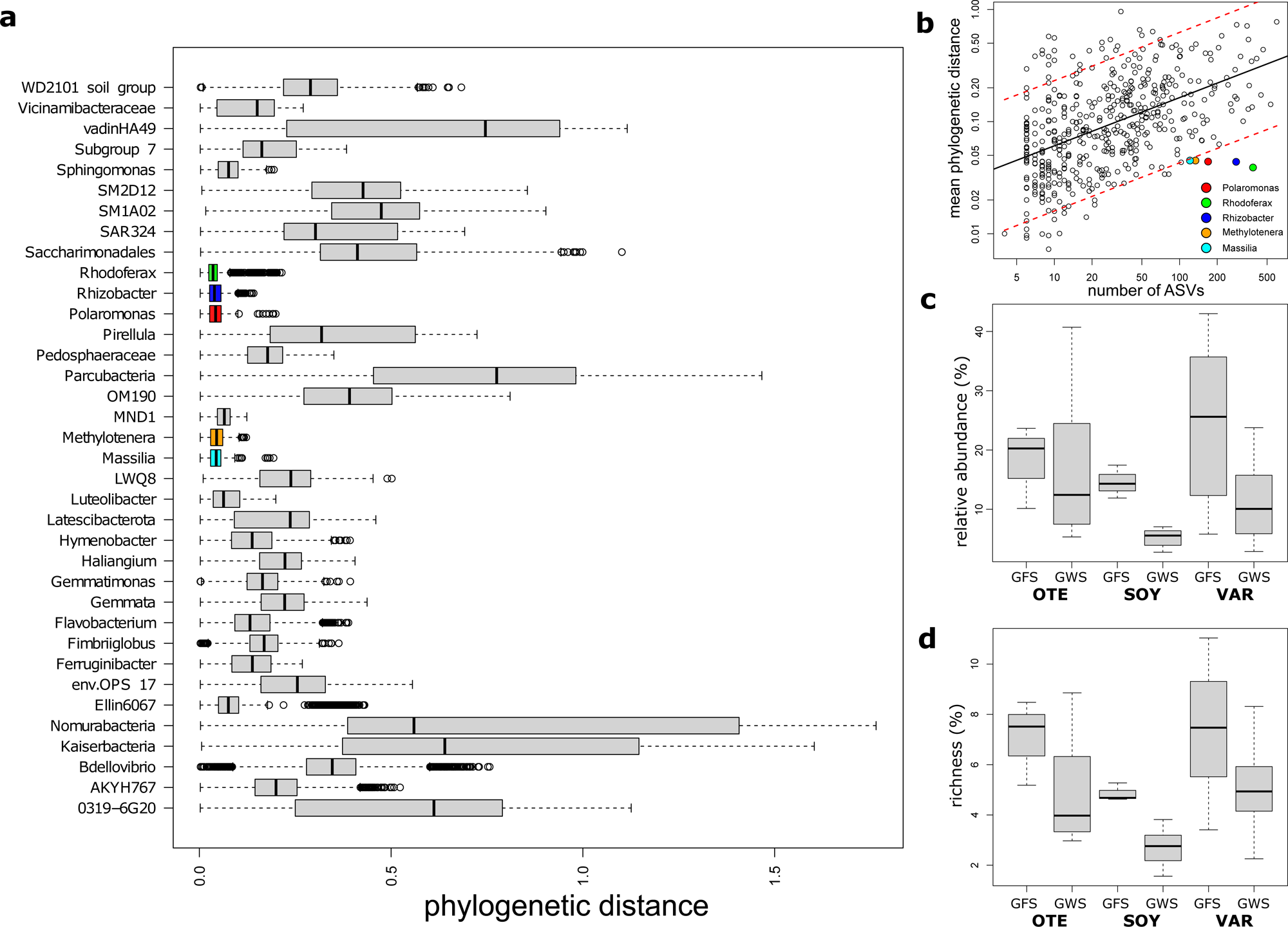
Microdiverse bacterial clades. Microdiverse bacterial genera are characterized by short pairwise phylogenetic distances among ASVs (a). Phylogenetic distances among ASVs with same genus-level taxonomy is influenced by the number of ASVs (b). Genera (circles) containing more ASVs generally have larger mean phylogenetic distances among members (regression line). Microdiverse genera fall below the 95% confidence intervals (red dashed lines) of this relationship, highlighting the high degree of phylogenetic clustering in these clades. Microdiverse clade members are particularly abundant in GFS (c) while they contribute disproportionately less to overall community richness (d).

We obtained 2732 high-quality bacterial MAGs and previous work unraveled the functional differences between MAGs found in GFS and GWS (Michoud et al., 2023). Here, we used MAGs primarily to identify phage-host interactions and screened them for their anti-phage arsenal. Nevertheless, MAGs, while only resolving a subset of the ASVs-resolved microbial diversity, reflected the composition of bacterial communities. Specifically, matching ASVs and MAGs with same genus-level classification, mean relative abundance of MAGs was well correlated with relative abundance of ASVs (Spearman Rho = 0.65, p<0.01).

### Viral community composition and diversity

Generally, little is known about viral biodiversity in extreme environments, particularly in mountain streams and rivers (Payne et al., 2020; Bekliz et al., 2022; Busi et al., 2022). Here, we identified 1452 high-quality (n=870) or complete (n=582) vMAGs. vMAG genome size varied considerably, with complete vMAG genome size averaging 47.8 kbp (range 2.6 to 373.3 kbp) and high-quality vMAG size averaging 70.4 kbp (range: 5.0 to 622.1 kbp). Screening for lysogeny-associated protein domains (e.g. integrases and recombinases), we found that 81.0% of complete vMAG genomes featured a virulent lifestyle (mean probability: 0.88 ± 0.1), whereas 19.0 % followed a temperate lifestyle (probability: 0.92 ± 0.11). Virulent vMAGs were typically less prevalent (found in 53.6 ± 1.4% of all samples) but more abundant (accounting for 16.3 ± 10.2% of vMAG relative abundance) than temperate vMAG (prevalence: 83.3 ± 1.6%, relative abundance: 2.9 ± 1.1%), without significant differences between GFS and GWS communities across the three floodplains. Contrary to Killing-the-winner and Piggyback-the-winner model expectations (Silveira and Rohwer, 2016; Chen et al., 2021), we found a constant ratio of temperate to virulent lifestyles (mean ratio: 0.37 ± 0.03) across a gradient of bacterial host abundances ranging from 1.8 × 10^5^ to 8.7 × 10^7^ cells per gram sediment.

Taxonomically, viral communities were dominated by dsDNA viruses classified as *Siphoviridae*, accounting on average for 35.6 and 37.5% of relative abundance in GFS and GWS, respectively (Figure 2 a,b). *Podoviridae* (16.5 and 17.7%) and *Myoviridae* (20.0 and 17.6%) were also prevalent and abundant in both stream types. Besides these members of *Caudovirales* (n=1165), another 25 viral families, including members of the ssDNA viral orders *Petitvirales* (n=70) and *Tubulavirales* (n=10) and of the eukaryotic-host infecting class of *Megaviricetes* (n=12) constituted the viral communities.

**Figure 2.**
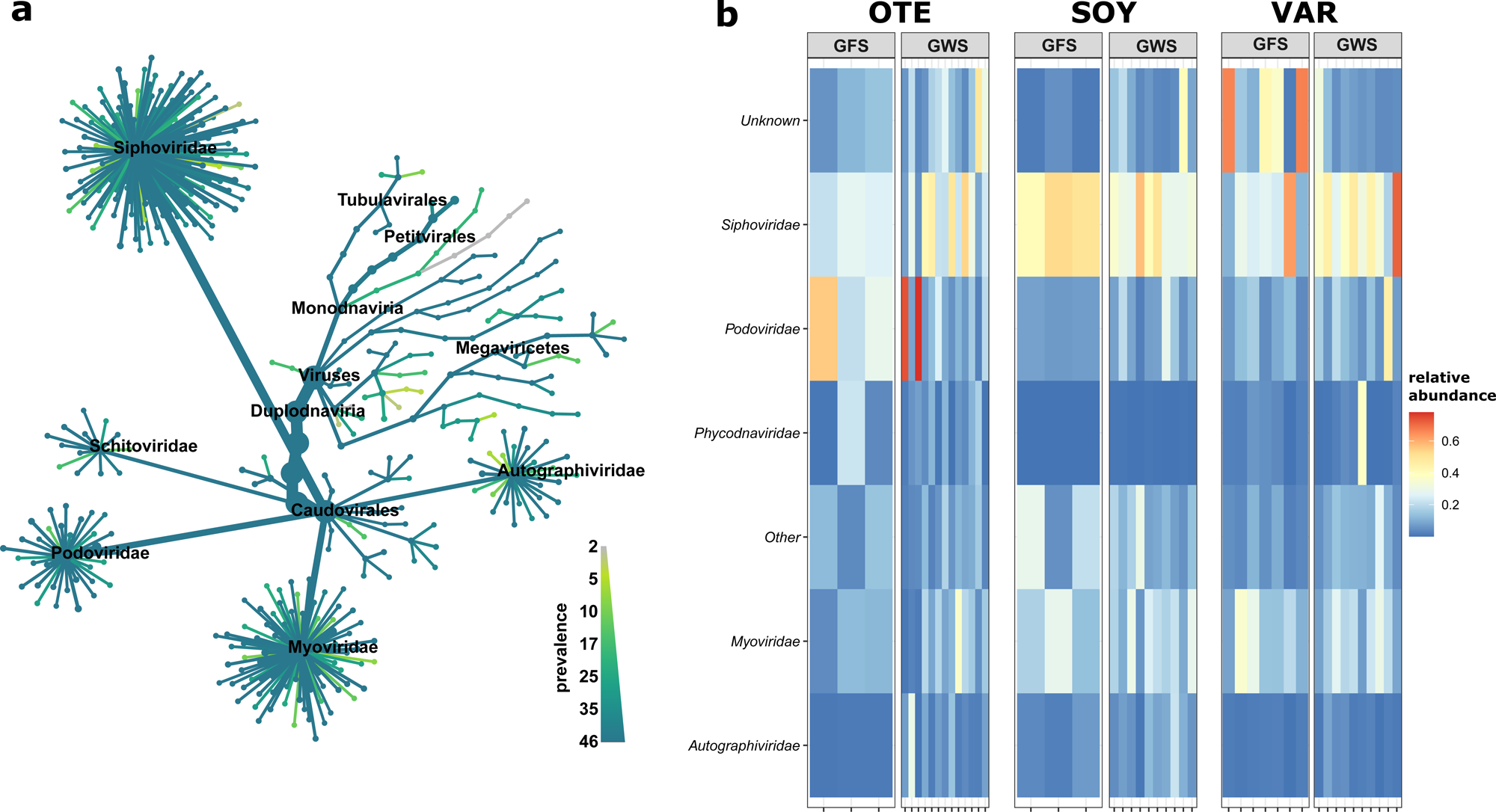
Viral taxonomic composition. vMAGs were classified mostly as members of Caudovirales, with high prevalence across most taxonomic groups (a). In terms of relative abundance, *Siphoviridae* vMAGs were particularly abundant in GWS, while unclassified vMAGs and vMAGs classified as *Podoviridae* and *Myoviridae* were more abundant in GFS (b).

Overall, vMAG prevalence was high with, on average, 1035.2 ± 50.8 vMAGs detected in each sample. Compared to the overall number of vMAGs (1452), this reflects the substantial overlap in terms of vMAG presence and absence across samples, irrespective of floodplain and stream type. Indeed, 455 vMAGs (31.3%) were detected in all samples, and vMAG prevalence, i.e. the percentage of samples in which a vMAG was detected, averaged 71.3 ± 31.7%. This large overlap in viral diversity was mainly driven by low-abundance vMAGs, i.e. vMAGs with a relative abundance of less than 0.005%. These low-abundance but highly prevalent vMAGs accounted on average for 25.7 ± 6.1% of all vMAGs detected in a sample, with no significant differences between floodplains (ANOVA, F=0.28, p=0.75) or stream types (ANOVA, F=0.28, p=0.60). Taken together, the high prevalence of mostly temperate phages suggests efficient dispersal mechanisms at the scale of Alpine proglacial streams. Given the topographic exposure and the relative isolation of proglacial environments, we suggest that temperate phages disperse efficiently embedded in the genomes of their prokaryotic hosts, whereas dispersal limitation is more important for virulent phages. This is relevant, as key environmental constraints for microbes in proglacial streams, such as repeated freeze-thawing, desiccation and exposure to UV radiation are similar during atmospheric transport. Hence, microbes particularly well adapted to these constraints may also efficiently disperse regionally, along with their temperate phages.

Across all floodplains and stream types, only few vMAGs were abundant (on average 7.8% of all vMAGs reached more than 0.1% of relative abundance in any given sample), yet, these abundant vMAGs accounted on average for 94.9% of total viral relative abundance. In contrast to the bacterial communities (Brandani et al., 2022), vMAG richness did not significantly differ between GFS and GWS (ANOVA, F=0.87, p=0.36) nor between the different floodplains (ANOVA, F=1.4, p= 0.25). On the one hand, metagenomics, as compared to viromics, may offer limited resolution to resolve rare phages. However, using metagenomics low viral richness has been linked to CRISPER-Cas abundance (Meaden et al., 2022), suggesting a trade-off exists between this adaptive bacterial immune response and phage diversity. We screened the bacterial antiviral defense arsenal and identified a total of 18,840 genes classified into 48 types (out of a total of 60 types included in DefenseFinder). Restriction-modification (ngenes=8,705), Cas (ngenes=4,483), abortive infection (ngenes=1,365), and the Septu defense system (ngenes=1,299) were the most commonly found defense types on bacterial MAGs. On average, 68.5 ± 7.8% of MAG relative abundance featured an antiviral defense system, with the above-mentioned types accounting for 57.2 ± 10.8% of overall relative abundance, highlighting the general importance of antiviral defense for bacterial stream biofilm communities. However, we did not find a significant relationship between viral diversity and Cas (Spearman Rho = 0.04, p=0.8) or any other defense system, nor differences in the relative abundance of different antiviral defense systems between stream types or floodplains. Taken together, the diverse virome and versatile antiviral defense arsenal in proglacial stream biofilms suggests a coevolutionary history of phages and biofilm-dwelling bacteria.

### Viral-bacterial community coupling

Bacterial community composition based on Bray-Curtis similarity obtained from ASV relative abundance significantly differed between floodplains (PERMANOVA, R^2^=0.08, p<0.001) and stream types (PERMANOVA, R^2^=0.03, p<0.001), with a significant interaction between floodplain and stream type (PERMANOVA, R^2^=0.05, p<0.001). Notably, bacterial communities from the Valsorey Glacier catchment were compositionally more distinct from communities of Otemma and ValRoseg, whereas particularly GWS sites of Otemma and Val Roseg were more similar (Figure 3a). This compositional similarity was mirrored by the viral community (Figure 3b), with significant differences between floodplains (PERMANOVA, R^2^=0.10, p<0.001) and stream types (PERMANOVA, R^2^=0.04, p<0.001), with a significant interaction term (PERMANOVA, R^2^=0.06, p<0.001). Superimposition of non-metric multidimensional scaling ordinations further substantiated the congruence between viral and bacterial communities (Procrustes correlation: 0.64, p=0.001, Figure 3c). The strength of these associations is comparable to earlier reports on the coupling of stream biofilm bacterial and viral communities (Bekliz et al., 2022) and suggests that host availability dominates over environmental conditions in shaping viral communities.

**Figure 3.**
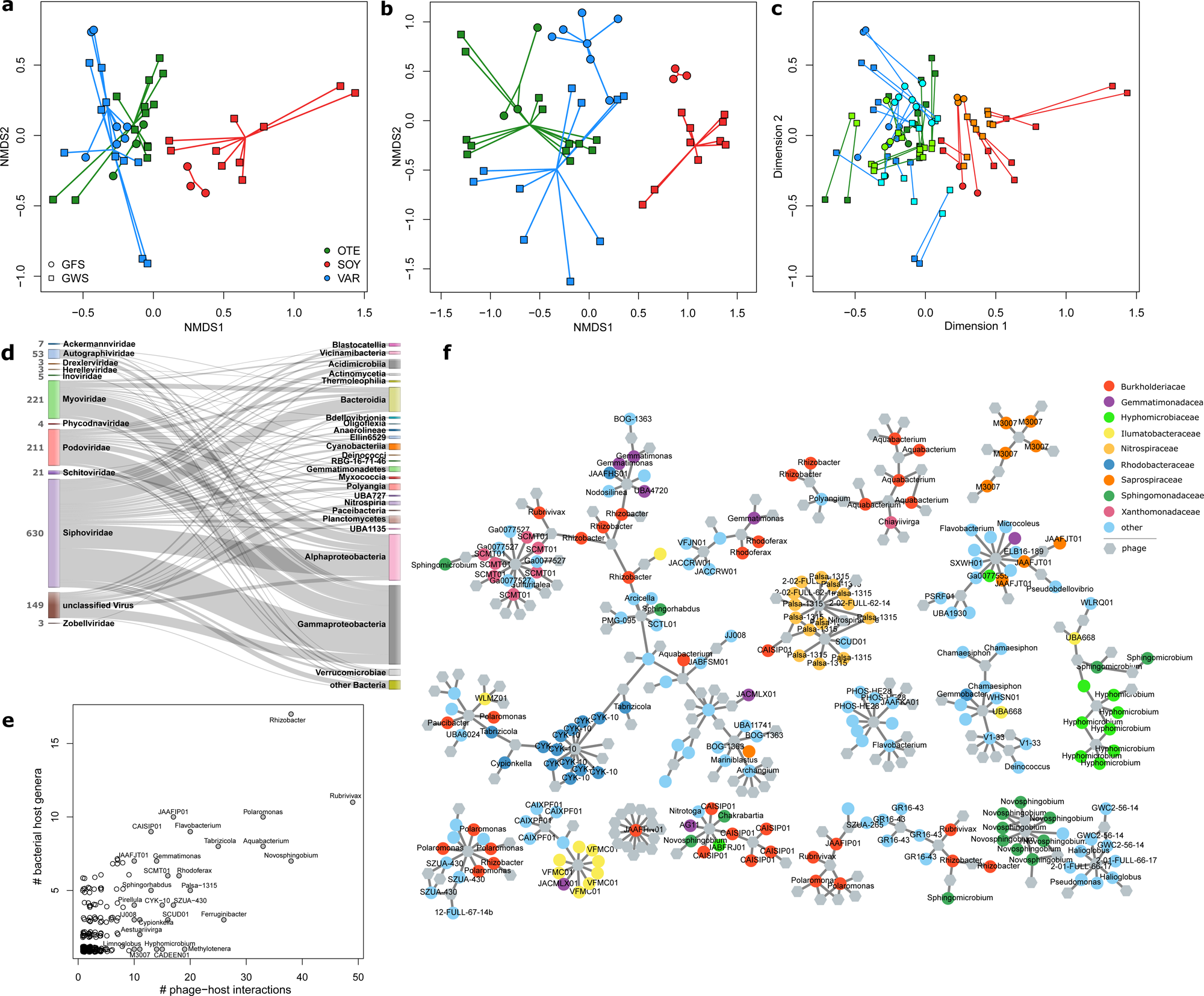
Phage-bacteria coupling. Non-metric multidimensional scaling ordinations based on vMAG relative abundance (a) and ASV relative abundance (b) reveal the compositional dissimilarities among floodplains and stream types. These differences were more pronounced in the bacterial communities, however, superpositioning of the two ordinations shows the significant coupling between phage and bacterial community similarity (c). Phage-host predictions summarized at broad taxonomic resolution (d) revealed that phages interact with diverse bacterial classes. Most phages interacted with a single or only few bacterial hosts (open circles, e). Shown are the number of phage-host interactions per bacterial genus (i.e. target genus) versus the total number of bacterial genera that phages infecting the target genus. The taxonomy for target genera with multiple phage host-interactions (filled circles) is displayed. Note that microdiverse genera, such as *Polaromonas* and *Rhizobacter* have many predicted phages, which also infect several other bacterial genera. Resolving putative interactions (grey lines) among individual vMAGs (hexagons) and bacterial MAGs (colored circles) illustrate the generally high phage-host specificity (i.e. phages tend to interact with bacteria of similar taxonomy) (e). The 17 largest sub-networks of the entire interaction network are shown.

We used phage-host predictions to further resolve the coupling between bacterial MAGs and vMAGs communities. PHIST predicted a total of 1435 phage-host interactions, which we compared to host-predictions obtained using iPHoP. Using a combination of the GTDB, the IMG isolate database and the GEM catalog as reference, iPHoP identified 283 host taxonomies for 227 vMAGs (i.e. some vMAGs obtained multiple iPHoP predictions). Despite the large viral and bacterial novelty in our samples, 53.8% of these predictions agreed with the PHIST-predicted host taxonomy at order level, suggesting robust k-mer frequency based phage-host predictions. Most PHIST-predicted interactions occurred between vMAGs classified as *Siphoviridae* and bacterial MAGs classified as Gammaproteobacteria (n=185), Alphaproteobacteria (n=137) and Bacteroidia (n=76; Figure 3d). vMAGs classified as *Podoviridae* or *Myoviridae* also interacted predominantly with proteobacterial MAGs (n=124 and n=31, respectively). Of the 953 bacterial MAGs with a predicted phage, particularly many phage-host interactions were identified involving MAGs classified as *Rubrivirax* (n=49), *Novosphingobium* (n=38), *Rhizobacter* (n=38), *Aquabacterium* (n=33) and *Polaromonas* (n=33)(Figure 3e). Similar to other work (Luo et al., 2022), the majority of phage-host interactions (84.2%) involved a single phage-host pair and only 6 phages had more than 10 predicted interactions with different MAGs (maxinteractions = 13). In line with generally high host specificity, vMAGs with multiple host interactions tended to interact with taxonomically similar MAGs (Figure 3e,f). For instance, 11 out of 13 interactions of a vMAG classified as *Lacusarxvirus* involved the alpha-proteobacterial CYK-10 genus. The other 2 interactions were predicted for MAGs classified as *Sphingomonas* (Alphaproteobacteria). Similarly, an unclassified vMAG interacted with 12 MAGs classified as *Nitrospiraceae*, an unclassified *Siphoviridae* vMAG interacted with 12 Gammaproteobacteria MAGs classified as *Xanthomonadales* (n=7) and *Burkholderiales* (n=5). Notably however, a vMAG classified as *Bertelyvirus* (*Siphoviridae*) interacted with 10 taxonomically distant MAGs classified as Gammaproteobacteria (n=5), Alphaproteobacteria (n=2), Bacteroidia (n=2) and Armatimonadota (n=1). Phages interacting with microdiverse clade members including *Rhizobacter*, *Polaromonas* and *Rhodoferax* also interact with several other bacterial genera (Figure 3e). For instance, of the 38 vMAGs putatively interacting with *Rhizobacter*, 22 vMAGs exclusively interacted with *Rhizobacter* whereas 16 vMAGs also interacted with different bacterial genera. However, this was not consistent for all microdiverse bacterial clades. All of the 19 phages predicted to interact with the microdiverse genus *Methylotenera*, exclusively interacted with *Methylotenera* MAGs.

While the limitations of *in silico* phage-host predictions need to be considered in this context (Brum and Sullivan, 2015; de Jonge et al., 2019), we report here a large number of putative phage-host interactions in stream biofilms, in line with reports from other extreme environments (Munson-McGee et al., 2018; Jarett et al., 2020). Moreover, the continuum of phage host range suggests that multiple eco-evolutionary processes shape bacteria-phage interactions in these communities (Weitz et al., 2013; de Jonge et al., 2019). On the one hand, trade-offs between virulence and host range (de Jonge et al., 2019) or resource limitation (Weitz et al., 2013) can explain narrow host ranges, whereas reduced dispersal and the dense packing of diverse bacteria have been associated with broader host ranges in biofilms (Hwang et al., 2023). In communities characterized by microdiversity, narrow host ranges may be beneficial given the increased encounter probability with phylogenetically closely related members of these clades. On the other hand, broad host range may increase the likelihood of exchanging beneficial genetic material, thus contributing to the longer-term success of bacterial clades.

### Auxiliary Metabolic Genes

In total, we detected 938 AMGs on 448 vMAGs, with a maximum of 42 AMGs on a single vMAG. Most AMGs were found on vMAGs classified as *Siphoviridae* (nvMAGs = 193), *Myoviridae* (nvMAGs = 111) and *Podoviridae* (nvMAGs = 61), broadly reflecting the taxonomic composition of the viral community and accounting for 35.3 ± 0.07% of vMAGs of these viral families. In contrast, AMGs were less common for other abundant vMAG families, including *Autographiviridae* (11 out of 55 vMAGs with AMGs), *Inoviridae* (1 out of 9 vMAGs) and particularly *Microviridae* (0 out of 70 vMAGs). In contrast to a recent analysis of Pearl River Estuary viromes^56^, we did not detect differences in relative prevalence of AMGs among virulent and temperate phages. Both viral lifestyles represented a similar number of AMGs when accounting for differences in lifestyle prevalence, and AMGs abundant among virulent phages were also abundant among temperate phages (Spearman Rho = 0.72, p<0.01).

AMGs encoded for a large functional diversity, with in total 166 different KEGG orthologues (KOs) present. Particularly numerous were AMGs encoding *dcm* (159 AMGs) which in prokaryotes can be involved in DNA methylation in restriction-modification systems (Pingoud et al., 2014). However, in phages, so-called orphan methyltransferases (i.e., without endonucleases of the prokaryotic restriction-modification system) can contribute to the protection of phage DNA against digestion (Heyerhoff et al., 2022). Other common AMGs were *queE, queD, queE and folE* genes (137 AMG), involved in the synthesis of the deazapurine nucleoside preQO, which also has been shown to protect viruses from restriction enzymes (Hutinet et al., 2019; Kieft et al., 2020). In contrast to pelagic systems (Kieft et al., 2021b; Heyerhoff et al., 2022) and despite the importance of cross-domain interactions in proglacial stream biofilms (Busi et al., 2022), however, AMGs encoding the core photosystem II proteins *psbA* and *psbD* were rare, with only 3 AMGs present across our entire dataset. The majority of AMGs encoded KOs involved in the metabolism of cofactors and vitamins (34.4%) including AMGs involved in folate biosynthesis, and the metabolisms of nicotinate/nicotinamide, and porphyrine/chlorophyll. AMGs involved in amino acid metabolism accounted for another 20.7% of all AMGs, including AMGs involved in cysteine and methionine metabolism. AMGs involved in carbohydrate metabolism (15.4%), glycan biosynthesis (8.4%) and sulfur metabolism (4.3%) were also commonly detected. Similar to marine sediments (Heyerhoff et al., 2022), *cysH*, which is involved in the synthesis of sulfite during assimilatory sulfate reduction, dominated AMGs involved in sulfur metabolism. *cysH* ranks among the globally conserved AMGs (Kieft et al., 2020) and has the potential to influence ecosystem-scale sulfur metabolism (Kieft et al., 2021a). This may be particularly relevant in GFS, where high turbidity and the absence of light fosters chemolithoautotrophic bacterial metabolisms. Chemolithoautotrophic bacteria can oxidize reduced sulfur to fix CO2 and to produce biomass. In the absence of photosynthetic organisms in GFS, chemolithoautotrophic carbon production may thus be an important ecosystem function and phage AMGs may play a pivotal role.

We did not detect significant differences in AMGs between viral communities from different floodplains or stream types (Figure 4a). Abundant AMG families, including the above-mentioned AMGs, tended to be present and abundant in most samples, whereas other AMG families were only sporadically present and often at low relative abundance. However, when coupling AMG with phage-host predictions (17.3% of AMGs were on vMAGs with a host prediction), we found that common and abundant AMGs were found on phages associated with specific bacterial community members (Figure 4b). For instance, 19 AMGs encoding folate biosynthesis were found on phages associated with 6 bacterial genera, including members of *Novosphingobium*, *Calothrix* and *Rhodobacter*. Similarly, AMGs encoding for cysteine and methionine metabolism were particularly prevalent among vMAGs infecting members of the bacterial genera *Polaromonas*, *Microcoleus*, *Ferruginibacter* and *Segetibacter*. Finally, using Principal Component Analysis with all AMGs for which phage-host predictions were available, we found distinct groups of bacterial genera to be host to phages with specific AMGs (Figure 4c). For example, the two microdiverse bacterial genera *Methylotenera* and *Rhizobacter*, together with *Lysobacter* and *Sandarakinorhabdus* formed a distinct group dominated by AMGs involved in porphyrine and chlorophyll metabolism. Another microdiverse bacterial genera, *Rhodoferax*, together with several other, non-microdiverse bacterial genera formed a distinct cluster based on AMGs involved in the metabolism of cysteine and methionine. AMGs involved in the biosynthesis of folate and lipopolysaccharides, explained clusters of bacterial genera involving members such as *Rhodobacter*, *Nitrospira*, *Novosphingobium* and *Calothrix* as well as a cluster composed of *Methylibium*, *Pedosphaera* and *Ideonella*. Interestingly, one of the most common bacterial genera in proglacial stream biofilms, *Polaromonas* did not associate with any of the clusters.

**Figure 4.**
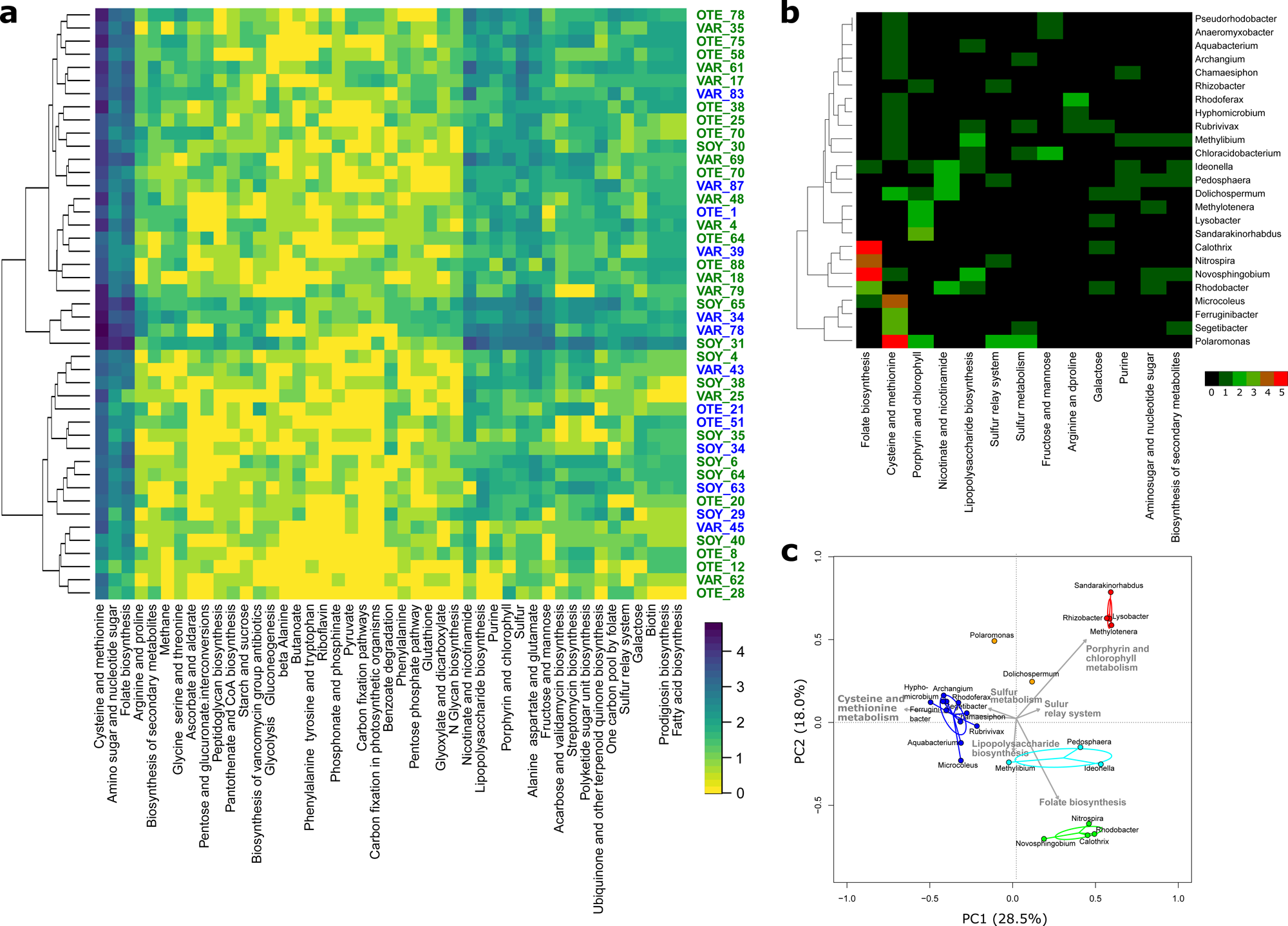
Auxiliary metabolic genes. A heatmap of the number of different AMGs found in the different samples highlights the prevalence of AMGs involved in the metabolism of cysteine and methionine, amino sugars and nucleotide sugar and folate biosynthesis (a). Despite the pronounced compositional difference between GFS and GWS, no differences in AMG composition between GFS (blue sample IDs) and GWS (green sample IDs) were apparent. Coupling AMGs with phage-host interactions revealed AMGs shared by phages putatively infecting bacterial MAGs (b). Considering all phage-host interactions and AMGs in a Principal Component Analysis, further highlighted the pronounced differences in phages carrying AMGs and infecting specific groups of bacteria (c). Note the distribution of microdiverse bacterial genera among these groups and the absence of the microdiverse bacterial genus *Massilia*.

Despite their compact genomes, phages regularly encode AMGs which can provide them a fitness advantage by augmenting or redirecting specific metabolic processes (Dion et al., 2020; Kieft et al., 2021b). This highlights the reciprocal benefits of phage-host interactions, which manifest during co-evolution (Koskella and Brockhurst, 2014). The selective environmental constraints in proglacial streams have persisted over geological time scales, shaping microbial diversity and putatively influencing phage-bacteria coevolution. Host specificity of AMGs has been previously reported (Luo et al., 2022) and here we report specific AMGs linked with microdiverse clades in proglacial stream biofilms. Arguably, these clades and their phages have shared a long evolutionary time in a selective environment, refining the AMGs library and putatively contributing to the eco-evolutionary success of both, the phages and their prokaryotic hosts. Microdiversity is an intrinsic property of many microbial communities and horizontal gene transfer can generate microdiversity (Larkin and Martiny, 2017). Phage-mediated HGT transfer may therefore contribute to microdiversification and narrow host range and AMG specificity suggests that in proglacial stream biofilms, virus-host interactions may foster ‘Maestro Microbes’ (Larkin and Martiny, 2017), which occupies niches by optimizing specific traits (in contrast to ‘Renaissance Microbes’ which acquire new traits, but see (Larkin and Martiny, 2017)).

## Conclusions

Generally, little is known about viral biodiversity in extreme environments, particularly however in streams and rivers draining mountain regions (Payne et al., 2020; Bekliz et al., 2022; Busi et al., 2022). Here, we expand the current knowledge of phage diversity and its role in shaping microbial communities in streams and rivers. Biofilms are thought to represent a physical barrier for viruses (Abedon, 2011), and our work unravels viral communities which are shaped by their bacterial host community composition. Extreme environmental conditions in proglacial streams lead to strong selective sorting of these host communities, ultimately resulting in communities dominated by few but well-adapted and microdiverse clades. Phages infecting members of these microdiverse clades carry AMGs which appear to be distinct and specific, suggesting that co-evolution between phages and their bacterial hosts shaped the potentially beneficial genetic repertoire of phages.

## Data availability

All sequence data are archived in the sequence read archive at NCBI under project number PRJNA808857.

## Author Contributions

HP, GM, SBB and TJB conceived the study. HP, GM and SBB performed bioinformatic and statistical analyses. All authors contributed to the writing of the article.

## Acknowledgements

We thank Jade Brandani, Nicola Deluigi, Kevin Casellini and Paraskevi Pramateftaki for assistance in the field and laboratory. We further thank Stylianos Fodelianakis, Tyler Kohler, Massimo Bourquin, Leila Ezzat and Martin Boutroux for fruitful discussions and feedback on the manuscript.

## References

Abedon, S. T. (2011). Bacteriophages and Biofilms: Ecology, Phage Therapy, Plaques. Nova Science.

Alneberg, J., Bjarnason, B. S., de Bruijn, I., Schirmer, M., Quick, J., Ijaz, U. Z., et al. (2014). Binning metagenomic contigs by coverage and composition. Nat Methods 11, 1144–1146. doi: 10.1038/nmeth.3103.

Bekliz, M., Pramateftaki, P., Battin, T. J., and Peter, H. (2022). Viral diversity is linked to bacterial community composition in alpine stream biofilms. ISME Communications. doi: 10.1038/s43705-022-00112-9.

Bolger, A. M., Lohse, M., and Usadel, B. (2014). Trimmomatic: a flexible trimmer for Illumina sequence data. Bioinformatics 30, 2114–2120. doi: 10.1093/bioinformatics/btu170.

Bolyen, E., Rideout, J. R., Dillon, M. R., Bokulich, N. A., Abnet, C. C., Al-Ghalith, G. A., et al. (2019). Reproducible, interactive, scalable and extensible microbiome data science using QIIME 2. Nat Biotechnol 37, 852–857. doi: 10.1038/s41587-019-0209-9.

Brandani, J., Peter, H., Busi, S. B., Kohler, T. J., Fodelianakis, S., Ezzat, L., et al. (2022). Spatial patterns of benthic biofilm diversity among streams draining proglacial floodplains. Frontiers in Microbiology 13.

Brandani, J., Peter, H., Fodelianakis, S., Kohler, T. J., Bourquin, M., Michoud, G., et al. (2023). Homogeneous Environmental Selection Structures the Bacterial Communities of Benthic Biofilms in Proglacial Floodplain Streams. Applied and Environmental Microbiology 89, e02010–22. doi: 10.1128/aem.02010-22.

Brum, J. R., and Sullivan, M. B. (2015). Rising to the challenge: accelerated pace of discovery transforms marine virology. Nat Rev Microbiol 13, 147–159. doi: 10.1038/nrmicro3404.

Busi, S. B., Bourquin, M., Fodelianakis, S., Michoud, G., Kohler, T. J., Peter, H., et al. (2022). Genomic and metabolic adaptations of biofilms to ecological windows of opportunity in glacier-fed streams. Nature Communications 13, 2168.

Busi, S. B., Pramateftaki, P., Brandani, J., Fodelianakis, S., Peter, H., Halder, R., et al. (2020). Optimised biomolecular extraction for metagenomic analysis of microbial biofilms from high-mountain streams. PeerJ 8, e9973. doi: 10.7717/peerj.9973.

Callahan, B. J., McMurdie, P. J., Rosen, M. J., Han, A. W., Johnson, A. J. A., and Holmes, S. P. (2016). DADA2: High-resolution sample inference from Illumina amplicon data. Nat Methods 13, 581–583. doi: 10.1038/nmeth.3869.

Chaumeil, P.-A., Mussig, A. J., Hugenholtz, P., and Parks, D. H. (2019). GTDB-Tk: a toolkit to classify genomes with the Genome Taxonomy Database. Bioinformatics 36, 1925–1927. doi: 10.1093/bioinformatics/btz848.

Chen, X., Weinbauer, M. G., Jiao, N., and Zhang, R. (2021). Revisiting marine lytic and lysogenic virus-host interactions: Kill-the-Winner and Piggyback-the-Winner. Science Bulletin 66, 871– 874. doi: 10.1016/j.scib.2020.12.014.

de Jonge, P. A., Nobrega, F. L., Brouns, S. J. J., and Dutilh, B. E. (2019). Molecular and Evolutionary Determinants of Bacteriophage Host Range. Trends Microbiol 27, 51–63. doi: 10.1016/j.tim.2018.08.006.

Dion, M. B., Oechslin, F., and Moineau, S. (2020). Phage diversity, genomics and phylogeny. Nat Rev Microbiol 18, 125–138. doi: 10.1038/s41579-019-0311-5.

Ezzat, L., Fodelianakis, S., Kohler, T. J., Bourquin, M., Brandani, J., Busi, S. B., et al. (2022). Benthic biofilms in glacier-fed streams from Scandinavia to the Himalayas host distinct bacterial communities compared with the streamwater. Applied and Environmental Microbiology 88, e00421–22.

Fodelianakis, S., Washburne, A. D., Bourquin, M., Pramateftaki, P., Kohler, T. J., Styllas, M., et al. (2022). Microdiversity characterizes prevalent phylogenetic clades in the glacier-fed stream microbiome. The ISME Journal 16, 666–675.

Foster, Z. S. L., Sharpton, T. J., and Grünwald, N. J. (2017). Metacoder: An R package for visualization and manipulation of community taxonomic diversity data. PLOS Computational Biology 13, e1005404. doi: 10.1371/journal.pcbi.1005404.

Freimann, R., Bürgmann, H., Findlay, S. E., and Robinson, C. T. (2013). Bacterial structures and ecosystem functions in glaciated floodplains: contemporary states and potential future shifts. ISME J 7, 2361–2373. doi: 10.1038/ismej.2013.114.

Gregory, A. C., Zayed, A. A., Conceição-Neto, N., Temperton, B., Bolduc, B., Alberti, A., et al. (2019). Marine DNA Viral Macro- and Microdiversity from Pole to Pole. Cell 177, 1109–1123.e14. doi: 10.1016/j.cell.2019.03.040.

Heyerhoff, B., Engelen, B., and Bunse, C. (2022). Auxiliary Metabolic Gene Functions in Pelagic and Benthic Viruses of the Baltic Sea. Front Microbiol 13, 863620. doi: 10.3389/fmicb.2022.863620.

Hockenberry, A. J., and Wilke, C. O. (2021). BACPHLIP: predicting bacteriophage lifestyle from conserved protein domains. PeerJ 9, e11396. doi: 10.7717/peerj.11396.

Hotaling, S., Hood, E., and Hamilton, T. L. (2017). Microbial ecology of mountain glacier ecosystems: biodiversity, ecological connections and implications of a warming climate. Environ Microbiol 19, 2935–2948. doi: 10.1111/1462-2920.13766.

Howard-Varona, C., Hargreaves, K. R., Abedon, S. T., and Sullivan, M. B. (2017). Lysogeny in nature: mechanisms, impact and ecology of temperate phages. ISME J 11, 1511–1520. doi: 10.1038/ismej.2017.16.

Hutinet, G., Kot, W., Cui, L., Hillebrand, R., Balamkundu, S., Gnanakalai, S., et al. (2019). 7-Deazaguanine modifications protect phage DNA from host restriction systems. Nat Commun 10, 5442. doi: 10.1038/s41467-019-13384-y.

Hwang, Y., Roux, S., Coclet, C., Krause, S. J. E., and Girguis, P. R. (2023). Viruses interact with hosts that span distantly related microbial domains in dense hydrothermal mats. Nat Microbiol, 1–12. doi: 10.1038/s41564-023-01347-5.

Jarett, J. K., Džunková, M., Schulz, F., Roux, S., Paez-Espino, D., Eloe-Fadrosh, E., et al. (2020). Insights into the dynamics between viruses and their hosts in a hot spring microbial mat. ISME J 14, 2527–2541. doi: 10.1038/s41396-020-0705-4.

Johansen, J., Plichta, D. R., Nissen, J. N., Jespersen, M. L., Shah, S. A., Deng, L., et al. (2022). Genome binning of viral entities from bulk metagenomics data. Nat Commun 13, 965. doi: 10.1038/s41467-022-28581-5.

Kang, D. D., Li, F., Kirton, E., Thomas, A., Egan, R., An, H., et al. (2019). MetaBAT 2: an adaptive binning algorithm for robust and efficient genome reconstruction from metagenome assemblies. PeerJ 7, e7359. doi: 10.7717/peerj.7359.

Kieft, K., Adams, A., Salamzade, R., Kalan, L., and Anantharaman, K. (2022). vRhyme enables binning of viral genomes from metagenomes. Nucleic Acids Research 50, e83. doi: 10.1093/nar/gkac341.

Kieft, K., Breister, A. M., Huss, P., Linz, A. M., Zanetakos, E., Zhou, Z., et al. (2021a). Virus-associated organosulfur metabolism in human and environmental systems. Cell Reports 36, 109471. doi: 10.1016/j.celrep.2021.109471.

Kieft, K., Zhou, Z., and Anantharaman, K. (2020). VIBRANT: automated recovery, annotation and curation of microbial viruses, and evaluation of viral community function from genomic sequences. Microbiome 8, 90. doi: 10.1186/s40168-020-00867-0.

Kieft, K., Zhou, Z., Anderson, R. E., Buchan, A., Campbell, B. J., Hallam, S. J., et al. (2021b). Ecology of inorganic sulfur auxiliary metabolism in widespread bacteriophages. Nat Commun 12, 3503. doi: 10.1038/s41467-021-23698-5.

Koskella, B., and Brockhurst, M. A. (2014). Bacteria-phage coevolution as a driver of ecological and evolutionary processes in microbial communities. FEMS Microbiol Rev 38, 916–931. doi: 10.1111/1574-6976.12072.

Krueger, F., James, F., Ewels, P., Afyounian, E., and Schuster-Boeckler, B. (2021). FelixKrueger/TrimGalore: v0.6.7 - DOI via Zenodo. doi: 10.5281/zenodo.5127899.

Larkin, A. A., and Martiny, A. C. (2017). Microdiversity shapes the traits, niche space, and biogeography of microbial taxa. Environ Microbiol Rep 9, 55–70. doi: 10.1111/1758-2229.12523.

Li, D., Liu, C.-M., Luo, R., Sadakane, K., and Lam, T.-W. (2015). MEGAHIT: an ultra-fast single-node solution for large and complex metagenomics assembly via succinct de Bruijn graph. Bioinformatics 31, 1674–1676. doi: 10.1093/bioinformatics/btv033.

Luo, X.-Q., Wang, P., Li, J.-L., Ahmad, M., Duan, L., Yin, L.-Z., et al. (2022). Viral community-wide auxiliary metabolic genes differ by lifestyles, habitats, and hosts. Microbiome 10, 190. doi: 10.1186/s40168-022-01384-y.

Meaden, S., Biswas, A., Arkhipova, K., Morales, S. E., Dutilh, B. E., Westra, E. R., et al. (2022). High viral abundance and low diversity are associated with increased CRISPR-Cas prevalence across microbial ecosystems. Current Biology 32, 220–227.e5. doi: 10.1016/j.cub.2021.10.038.

Menzel, P., Ng, K. L., and Krogh, A. (2016). Fast and sensitive taxonomic classification for metagenomics with Kaiju. Nat Commun 7, 11257. doi: 10.1038/ncomms11257.

Michoud, G., Kohler, T. J., Peter, H., Brandani, J., Busi, S. B., and Battin, T. J. (2023). Unexpected functional diversity of stream biofilms within and across proglacial floodplains despite close spatial proximity. Limnology and Oceanography. doi: 10.1002/lno.12415.

Munson-McGee, J. H., Peng, S., Dewerff, S., Stepanauskas, R., Whitaker, R. J., Weitz, J. S., et al. (2018). A virus or more in (nearly) every cell: ubiquitous networks of virus-host interactions in extreme environments. ISME J 12, 1706–1714. doi: 10.1038/s41396-018-0071-7.

Nayfach, S., Camargo, A. P., Schulz, F., Eloe-Fadrosh, E., Roux, S., and Kyrpides, N. C. (2021). CheckV assesses the quality and completeness of metagenome-assembled viral genomes. Nat Biotechnol 39, 578–585. doi: 10.1038/s41587-020-00774-7.

Needham, D. M., Sachdeva, R., and Fuhrman, J. A. (2017). Ecological dynamics and co-occurrence among marine phytoplankton, bacteria and myoviruses shows microdiversity matters. ISME J 11, 1614–1629. doi: 10.1038/ismej.2017.29.

Nissen, J. N., Johansen, J., Allesøe, R. L., Sønderby, C. K., Armenteros, J. J. A., Grønbech, C. H., et al. (2021). Improved metagenome binning and assembly using deep variational autoencoders. Nat Biotechnol 39, 555–560. doi: 10.1038/s41587-020-00777-4.

Obeng, N., Pratama, A. A., and Elsas, J. D. van (2016). The Significance of Mutualistic Phages for Bacterial Ecology and Evolution. Trends Microbiol 24, 440–449. doi: 10.1016/j.tim.2015.12.009.

Oksanen, J., Blanchet, F. G., Kindt, R., Legendre, P., Minchin, P. R., O’hara, R. B., et al. (2013). Community ecology package. R package version 2.

Olm, M. R., Brown, C. T., Brooks, B., and Banfield, J. F. (2017). dRep: a tool for fast and accurate genomic comparisons that enables improved genome recovery from metagenomes through de-replication. ISME J 11, 2864–2868. doi: 10.1038/ismej.2017.126.

Parks, D. H., Imelfort, M., Skennerton, C. T., Hugenholtz, P., and Tyson, G. W. (2015). CheckM: assessing the quality of microbial genomes recovered from isolates, single cells, and metagenomes. Genome Res 25, 1043–1055. doi: 10.1101/gr.186072.114.

Payne, A. T., Davidson, A. J., Kan, J., Peipoch, M., Bier, R., and Williamson, K. (2020). Widespread cryptic viral infections in lotic biofilms. Biofilm 2, 100016.

Pingoud, A., Wilson, G. G., and Wende, W. (2014). Type II restriction endonucleases--a historical perspective and more. Nucleic Acids Res 42, 7489–7527. doi: 10.1093/nar/gku447.

Ren, J., Song, K., Deng, C., Ahlgren, N. A., Fuhrman, J. A., Li, Y., et al. (2020). Identifying viruses from metagenomic data using deep learning. Quant Biol 8, 64–77. doi: 10.1007/s40484-019-0187-4.

Roux, S., Brum, J. R., Dutilh, B. E., Sunagawa, S., Duhaime, M. B., Loy, A., et al. (2016). Ecogenomics and potential biogeochemical impacts of globally abundant ocean viruses. Nature 537, 689– 693.

Roux, S., Camargo, A. P., Coutinho, F. H., Dabdoub, S. M., Dutilh, B. E., Nayfach, S., et al. (2023). iPHoP: An integrated machine learning framework to maximize host prediction for metagenome-derived viruses of archaea and bacteria. PLOS Biology 21, e3002083. doi: 10.1371/journal.pbio.3002083.

Schwartz, D. A., Lehmkuhl, B. K., and Lennon, J. T. (2022). Phage-encoded sigma factors alter bacterial dormancy. Msphere 7, e00297–22.

Shannon, P., Markiel, A., Ozier, O., Baliga, N. S., Wang, J. T., Ramage, D., et al. (2003). Cytoscape: a software environment for integrated models of biomolecular interaction networks. Genome Res 13, 2498–2504. doi: 10.1101/gr.1239303.

Shkoporov, A. N., Turkington, C. J., and Hill, C. (2022). Mutualistic interplay between bacteriophages and bacteria in the human gut. Nat Rev Microbiol 20, 737–749. doi: 10.1038/s41579-022-00755-4.

Sieber, C. M. K., Probst, A. J., Sharrar, A., Thomas, B. C., Hess, M., Tringe, S. G., et al. (2018). Recovery of genomes from metagenomes via a dereplication, aggregation and scoring strategy. Nat Microbiol 3, 836–843. doi: 10.1038/s41564-018-0171-1.

Silveira, C. B., and Rohwer, F. L. (2016). Piggyback-the-Winner in host-associated microbial communities. npj Biofilms Microbiomes 2, 1–5. doi: 10.1038/npjbiofilms.2016.10.

Stamatakis, A. (2014). RAxML version 8: a tool for phylogenetic analysis and post-analysis of large phylogenies. Bioinformatics 30, 1312–1313. doi: 10.1093/bioinformatics/btu033.

Suttle, C. (1994). The Significance of Viruses to Mortality in Aquatic Microbial Communities. Microbial ecology 28, 237–43. doi: 10.1007/BF00166813.

Suttle, C. A. (2007). Marine viruses — major players in the global ecosystem. Nat Rev Microbiol 5, 801–812. doi: 10.1038/nrmicro1750.

Tesson, F., Hervé, A., Mordret, E., Touchon, M., d’Humières, C., Cury, J., et al. (2022). Systematic and quantitative view of the antiviral arsenal of prokaryotes. Nat Commun 13, 2561. doi: 10.1038/s41467-022-30269-9.

Weitz, J. S., Poisot, T., Meyer, J. R., Flores, C. O., Valverde, S., Sullivan, M. B., et al. (2013). Phage-bacteria infection networks. Trends Microbiol 21, 82–91. doi: 10.1016/j.tim.2012.11.003.

Zielezinski, A., Deorowicz, S., and Gudyś, A. (2022). PHIST: fast and accurate prediction of prokaryotic hosts from metagenomic viral sequences. Bioinformatics 38, 1447–1449. doi: 10.1093/bioinformatics/btab837.

Zimmerman, A. E., Howard-Varona, C., Needham, D. M., John, S. G., Worden, A. Z., Sullivan, M. B., et al. (2020). Metabolic and biogeochemical consequences of viral infection in aquatic ecosystems. Nat Rev Microbiol 18, 21–34. doi: 10.1038/s41579-019-0270-x.

